# RORγt-Expressing Pathogenic CD4^+^T Cells Cause Brain Inflammation During Chronic Colitis

**DOI:** 10.1101/2021.09.01.458634

**Authors:** Michel Edwar Mickael, Suniti Bhaumik, Ayanabha Chakraborti, Alan Umfress, Thomas van Groen, Matthew Macaluso, John Totenhagen, Anna G Sorace, James A Bibb, David G Standaert, Rajatava Basu

## Abstract

Neurobehavioral disorders and brain abnormalities have been extensively reported in both Crohn’s Disease (CD) and Ulcerative Colitis (UC) patients. However, the mechanism causing neuropathological disorders in inflammatory bowel disease (IBD) patients remains unknown. Studies have linked the Th17 subset of CD4^+^T cells to brain diseases associated with neuroinflammation and cognitive impairment, including multiple sclerosis (MS), ischemic brain injury and Alzheimer’s disease. To better understand how CD4^+^T lymphocytes, contribute to brain pathology in chronic intestinal inflammation, we investigated the development of brain inflammation in the T cell transfer model of chronic colitis. Our findings demonstrate that CD4^+^T cells infiltrate the brain of colitic *Rag1*^-/-^ mice in proportional levels to colitis severity. Colitic mice developed hypothalamic astrogliosis that correlated with neurobehavioral disorders. Moreover, the brain-infiltrating CD4^+^T cells expressed Th17 cell transcription factor RORγt and displayed a pathogenic Th17 cellular phenotype similar to colonic Th17 cells. Adoptive transfer of RORγt-deficient naïve CD4^+^T cells failed to cause brain inflammation and neurobehavioral disorders in *Rag1*^-/-^ recipients, with significantly less brain infiltration of CD4^+^T cells. These findings suggest that pathogenic RORγt^+^CD4^+^T cells that aggravate colitis migrate preferentially into the brain, contributing to brain inflammation and neurobehavioral disorders, thereby linking colitis severity to neuroinflammation.

## Introduction

Growing evidence suggests that disturbance in the gut-brain axis in inflammatory bowel disease (IBD) is associated with neuropathological conditions. Both symptomatic and asymptomatic CNS abnormalities as well as neurobehavioral disorders like anxiety and depression have been reported extensively in both Crohn’s Disease (CD) and Ulcerative Colitis (UC) patients (1-10). In both types of IBD, a high degree of correlation with several neurodegenerative disorders such as multiple sclerosis (MS)-like conditions, dementia, and Parkinson disease have been reported (11-14). Demyelinating disorder is observed occur more commonly among patients with IBD compared to non-IBD patients (11, 15). IBD patients can present various manifestations, including vascular injuries, inflammation of blood vessels in the brain, and increased neural infections (10). Particularly, research show that 4 in 10 people with IBD experience feelings of anxiety and/or depression and IBD patients were significantly more likely to have a lifetime diagnosis of major anxiety and depression (8, 9). The limbic system including the hippocampus and the hypothalamus constitute one of the main areas of the brain that have been implicated in depression and anxiety disorders related to colitis. Decreased hippocampal neurogenesis has been demonstrated in colitic mice (16). The hypothalamus is implicated in emotion processing and plays a major role in depression, anxiety, and pathology of mood disorders (17-19). It has been demonstrated that psychological stress acting through the hypothalamic-pituitary-adrenal (HPA) axis increases intestinal motility in IBD patients, which causes local inflammation in the gastro-intestinal (GI) tract and worsens IBD (7, 20, 21). In turn, gut inflammation in IBD patients is believed to activate peripheral inflammatory responses leading to the development of mood disorders associated with hypothalamus (22).

The cause-and-effect relationship between IBD and neurological comorbidity is controversial (10, 20). Researchers have previously demonstrated that intestinal inflammation can influence brain activity and psychological behavior in preclinical models (20). Studies of animal models with colonic inflammation have shown inflammatory changes in the brain (23-25). Despite these observations, little is known about the main mechanistic factors behind these phenomena (1, 2, 5, 6). It has yet to be established whether hyperactive immune response associated with inflammation of the gut triggers brain inflammation that results in neuropathological disorders during IBD. Possible explanations including vasculitis, thromboembolism and malnutrition have been proposed (10). Although it is known that proinflammatory CD4^+^T cells contribute to the pathogenesis of both IBD and MS (26, 27), whether IBD can trigger infiltration of gut-derived proinflammatory CD4^+^T cells into the brain has not been investigated. New studies have demonstrated roles of a broad range of immune cells, including innate lymphoid cells and T cells, in regulating inflammation and neurological diseases in the brain (28). Moreover, IL-17 and IFNγ-producing autoreactive CD4^+^ T cells are the main pathogenic populations that infiltrate the central nervous system (CNS) and determine the clinical course of autoimmune disease of the brain (29). Among the different subsets of CD4^+^T cells, Th17 cells are more effective inducers of neuroinflammation compared to Th1 cells as they cross blood brain barrier (BBB) more effectively and preferentially target the astrocytes (30-32). However, their role in modulating gut-brain axis during IBD is not yet known.

Due to the unusual association of IBD with neuropathological comorbidity, we asked 2 central questions: (1) Does chronic colitis cause inflammation in any part of the brain leading to neuropathological disorders? (2) How does chronic colitis cause brain inflammation? To understand whether neuropathological conditions during IBD represents a simple association or a causation of IBD, we investigated the development of brain inflammation in a mouse model of chronic colitis. We found a significant correlation between gastrointestinal inflammation and pro-inflammatory response in the brain. Moreover, this link between peripheral and CNS inflammation appeared to be causally linked to deleterious effects on brain neurobehavioral function. Therefore, these findings suggest that gut-derived CD4^+^T cells trigger inflammation in the CNS that contributes to the pathogenesis of anxiety and depression in IBD patients.

## Results

### Result 1: T cell mediated chronic colitis show CD4^+^T cell infiltration in the brain that is proportional to colitis severity

IBD is often associated with neurodegenerative disorders and neuropsychiatric manifestations such as depression, fatigue, and anxiety (1, 10, 33-35). To understand whether IBD drives brain inflammation, we investigated the development of brain inflammation in the adoptive T cell transfer model of chronic colitis in *Rag1*^-/-^ mice which lack T and B cells. While the brains of normal un-transferred *Rag1*^*-/-*^ mice are devoid of CD4^+^T cells, a substantial population of CD4^+^T cells accumulates in the brain of colitic *Rag1*^-/-^ mice starting from the onset of colitis at 4 wk post CD45RB^hi^ CD4^+^T cell transfer. At 8wk post-transfer when colitis becomes severe,∼15% of the total live brain mononuclear cells were CD4^+^T cells (**Fig.1 A, B**). Between 4 and 8 wk post-transfer, there was >5 -fold increase in the number of brain-infiltrating CD4^+^T cells that was proportional to the severity of colitis (**Fig. 1B, C**). We found a significant correlation, between the increase in the number of colonic CD4^+^T cells and brain infiltrating CD4^+^T cells as severity of disease progressed over time (**Fig. 1D**). Among several organs analyzed in colitic mice, the number of CD4^+^T cell increased by >20-fold only in the gut and in the brain during the peak phase of colitis (4 - 8 wk) whereas all other organs showed little increase (<2-fold) in the number of CD4^+^T cells between the onset and development of severe colitis (**Fig. 1E**). This indicated that the number of CD4^+^T cell in the brain increases proportionally to the expansion of effector CD4^+^T cells in the gut. To exclude the possibility that the observed phenomenon is not specific to colitis induction in lymphopenic *Rag1*^-/-^ mice, we analyzed the extent of CD4^+^T cell infiltration in a chronic model of Dextran Sodium Sulfate-induced colitis (cDSS) in normal lympho-sufficient B6 mice. Similar to the observation in the adoptive transfer model of colitis in *Rag1*^*-/-*^ mice, we found marked infiltration of CD4^+^T cells in the brain of DSS-induced colitic mice where the frequency of brain-infiltrating CD4^+^T cells was directly proportional to colitis severity that increased in each cycle of DSS treatment (**Fig. S1**). After 3 cycles of DSS treatment, there was ∼25-fold increase in the frequency of brain infiltrating CD4^+^T cells and a ∼300-fold increase in the total number of CD3^+^CD4^+^T cells in the brain compared to vehicle-treated control (0.3-0.5% CD4^+^T cells or absolute number CD4^+^ T cells were ∼350 in age matched normal WT mice). Together, our data indicated that CD4^+^T cells infiltrate the brain of mice during colitis and number of infiltrating CD4^+^T cell into the brain is proportional to progression in disease severity.

**Figure 1:**
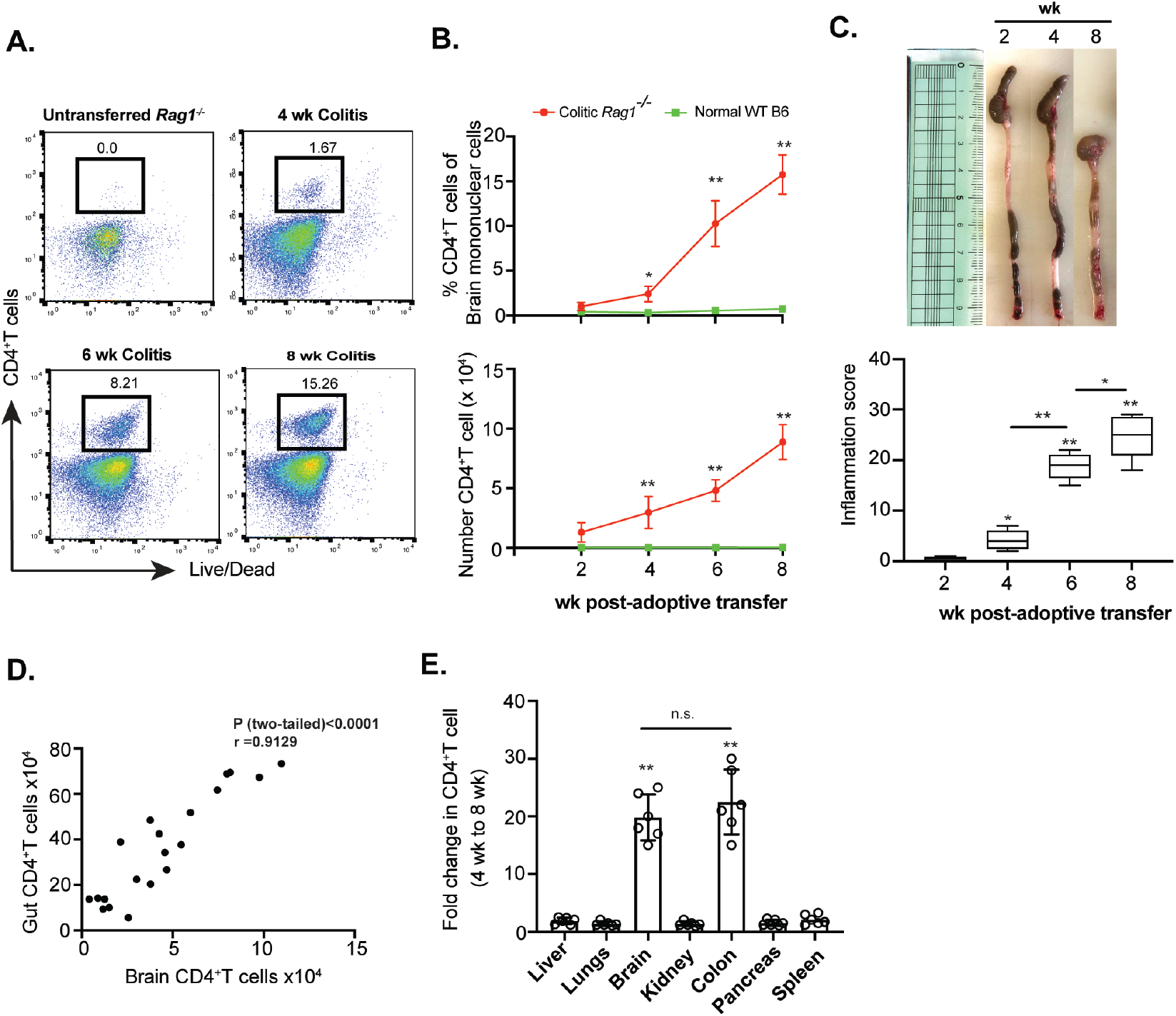
CD4^+^T cells infiltrate into the brain during chronic colitis. **(A)** Sorted naïve CD4^+^Tcells (4×10^5^/mouse i.p.) from normal B6 mice were adoptively transferred to *Rag1*^*-/-*^ mice for colitis induction and brain-infiltrating CD4^+^T cell frequency was analyzed by flow cytometry at different time points post-transfer compared to normal un-transferred *Rag1*^*-/-*^ mice represented by FACS plots. **(B)** Time kinetics of percentage (top) and total numbers (bottom) of retrieved brain CD4^+^T cells from CD45RB^hi^ CD4^+^T cells recipient colitic *Rag1*^-/-^ mice compared to age and sex-matched normal *Rag1*^*-/-*^ mice at different time points (n=9). **(C)** Colonic length (top) and intestinal inflammation score (bottom) of *Rag1*^*-/-*^ recipient mice at different time points post-adoptive transfer of CD45RB^hi^ CD4^+^ T cells (n=8). **(D)** Scatter plot showing positive correlation between gut-derived and brain-derived CD4^+^T cells of colitic mice as analyzed between 2 wk and 8 wk post-adoptive transfer (n=20). **(E)** Fold change in number of CD4^+^T cells retrieved from different organs between 4 wk and 8 wk post-transfer in naïve CD4^+^ T cells recipient *Rag1*^*-/-*^ mice showing significant fold-increase in colonic and brain-derived CD4^+^T cell number. Data are shown as mean ± SEM. Data are representative of 3 independent experiments. *P* values, one-way ANOVA followed by Tukey’s post hoc test (B, C, E); Pearson Correlation coefficient with indicated R and two-tailed P value <0.0001 (D). *p<0.01; #p<0.005; **p<0.0001.

### Result 2. Development of chronic colitis is associated with anxiety and depression-like neurobehavioral disorders

In behavioral neuroscience, quantitative responses to stressful environments such as those presented in the which elevated plus maze (EPM), social interaction (sociability), and light-dark tests may be used to approximate assessment of emotional state. Aversion to stressful components of these tests has been suggested to evidence anxiodepressive-like behavior in rodents (36-39). To understand whether brain inflammation correlated with neurobehavioral abnormality, colitic mice were tested in these paradigms (40).

Colitic *Rag1*^-/-^ mice adoptively transferred with naïve CD4^+^T cells exhibited increased anxiodepressive-like behavior compared to controls in the EPM, light-dark test, and social interaction test (**Fig. 2A, 2B**). Interestingly, despite these deficiencies, the performance of colitic mice was comparable to normal control mice (un-transferred *Rag1*^-/-^) with regard to learning and memory. Specifically, the ability to remember and identify novel objects was not affected in colitic mice as evident from novel object recognition test. Also, there was no effect on aversive memory formation or working memory as inducted by assessments in the fear conditioning and Y-maze tests, respectively. Similarly, the colitic *Rag1*^-/-^ mice did not show a significant difference in missteps compared to control mice in the horizontal ladder test suggesting the absence of any defect in their motor function. Thus, colitic mice showed acute anxiety and depression-like behavior with no apparent deficits in cognition, working memory, spatial memory, cerebellum function, and muscle coordination. These data are consistent with the correlations between gastrointestinal inflammation and neuropsychiatric conditions observed in humans.

**Figure 2:**
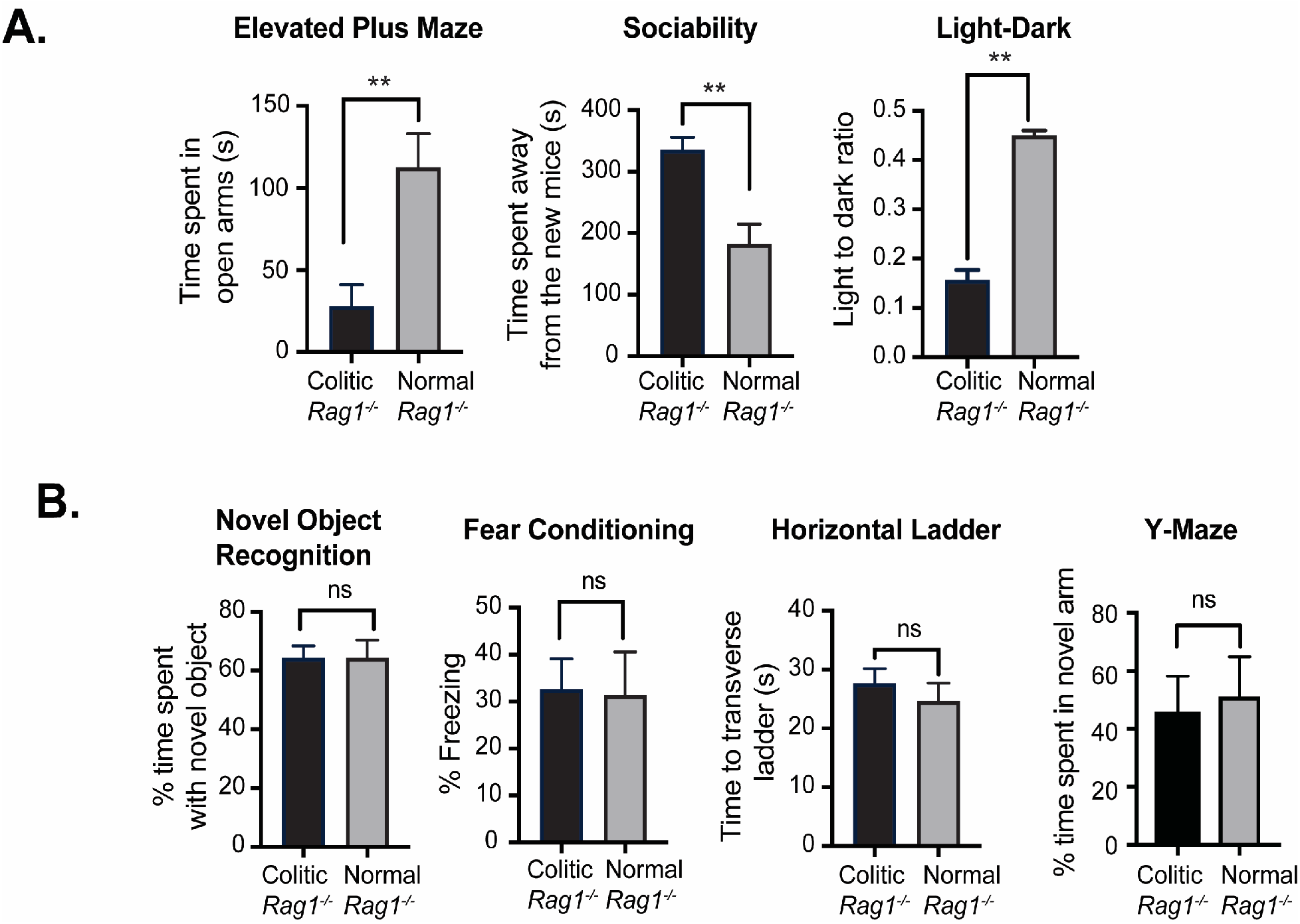
Colitic mice anxiety and depression-like neurobehavioral disorders but not memory or motor deficiency. (A) Analysis of comparative neurobehavioral study by Elevated Maze Test, Sociability, Light Dark test. (B) Analysis of Novel Object Recognition, Fear Conditioning, Horizontal Ladder and Y-Maze test between normal *Rag1*^*-/-*^ mice and CD45RB^hi^ CD4^+^T cells recipient colitic *Rag1*^-/-^ mice at 10 wk post-transfer (n= 6 mice/group). Data are shown as mean ± SEM. Data are representative of 2 independent experiments. *P* values, two-tailed unpaired Student *t*-test; **p<0.0001; n.s.= not significant.

### Result 3. Colitic mice show brain inflammation with CD4^+^T cell accumulation and astrogliosis in the hypothalamus

Dysregulation in the hypothalamus has been found to play a critical role in stress and depression during IBD (41). Targeting certain neurons in the hypothalamus rather than the whole brain potentially provides a more effective treatment for anxiety than conventional therapy (19). Neurobehavioral manifestations including anxiety and stress have been noted for many years in IBD patients (33, 42). As gastrointestinal disorders and hypothalamus-pituitary-adrenal (HPA) axis dysfunction are frequently observed in patients with major anxiety and depression, we analyzed the neurobehavioral outcome of the hypothalamus (43). Recently, it has been reported that the hippocampus neurogenesis is significantly affected in colitis. This alteration in neurodegeneration has been linked to anxiety-like behavior in a colitic mouse model(16). Thus, we specifically concentrated on these two main brain regions in our subsequent investigations. As the behavioral studies suggested that the colitis impacted anxiodepressive-like parameters but not motor or memory functions, we focused on the limbic system circuitry controlling these emotions. While the hippocampus can also contribute to depression and anxiety, as well as food intake, we found a substantial CD4^+^ T cell accumulation in the hypothalamus. However, significantly lower number of CD4^+^T cells migrated to the hippocampus compared to the hypothalamus (**Fig.3A, C**). Interestingly, diffusion-weighted MRI (DWI) of 10 wk post-transferred *Rag1*^-/-^ colitic mice revealed a small hyperintense signal in the hypothalamus including the preoptic region near the AV3V (**Fig.3B**).

In the CNS, astrocytes respond rapidly to any type of neurologic insults through a process called astrogliosis by the up-regulation of intermediate filament, glial fibrillary acidic protein (GFAP) indicating astroglial activation (44). As colitic *Rag1*^-/-^ mice showed a lesion-like structure in the hypothalamic region with accumulation of CD4^+^T cells, we examined cross-sections of the brain of colitic mice for accumulation of reactive astrocytes at 8 wk post transfer. In colitic mice, a marked expansion of the astrocytes occurred in the periventricular hypothalamic region, with pronounced up-regulation of GFAP, cellular hypertrophy and dispersed astrocyte proliferation, which are classical signs of ongoing inflammation in the brain but not in the hippocampus area (**Fig. 3D, E**). Astrocytes have long been related to demyelinating diseases of the CNS and as reactive components within and surrounding demyelinated lesions in MS (45). Analysis of the hypothalamic region of colitic mouse brain also revealed significant demyelination (**Fig. S2**). The appearance of demyelination in human IBD patients is controversial. The vast majority of patients with UC present a form of neurobehavioral problems, including anxiety and depression (1-10). Nevertheless, it has been claimed that only 3% of patients suffering from IBD manifest visible demyelinating lesions during the course of the disease(11, 15). Additionally, it has been shown that 42-46 % of IBD have small white matter lesions, compared with 16% in controls (1). Interestingly, astrogliosis and demyelination have been described in monkeys suffering from UC (13). Together, the colitic mice show a high degree of astrogliosis, demyelination and lesion-like development in the hypothalamic area that are associated with neurobehavioral disorders. Therefore, it is likely that astrogliosis with signs of demyelination are critical steps in the initiation of T cell-mediated pathology in the CNS as well as indicative of ongoing inflammation in the brain during chronic colitis.

**Figure 3:**
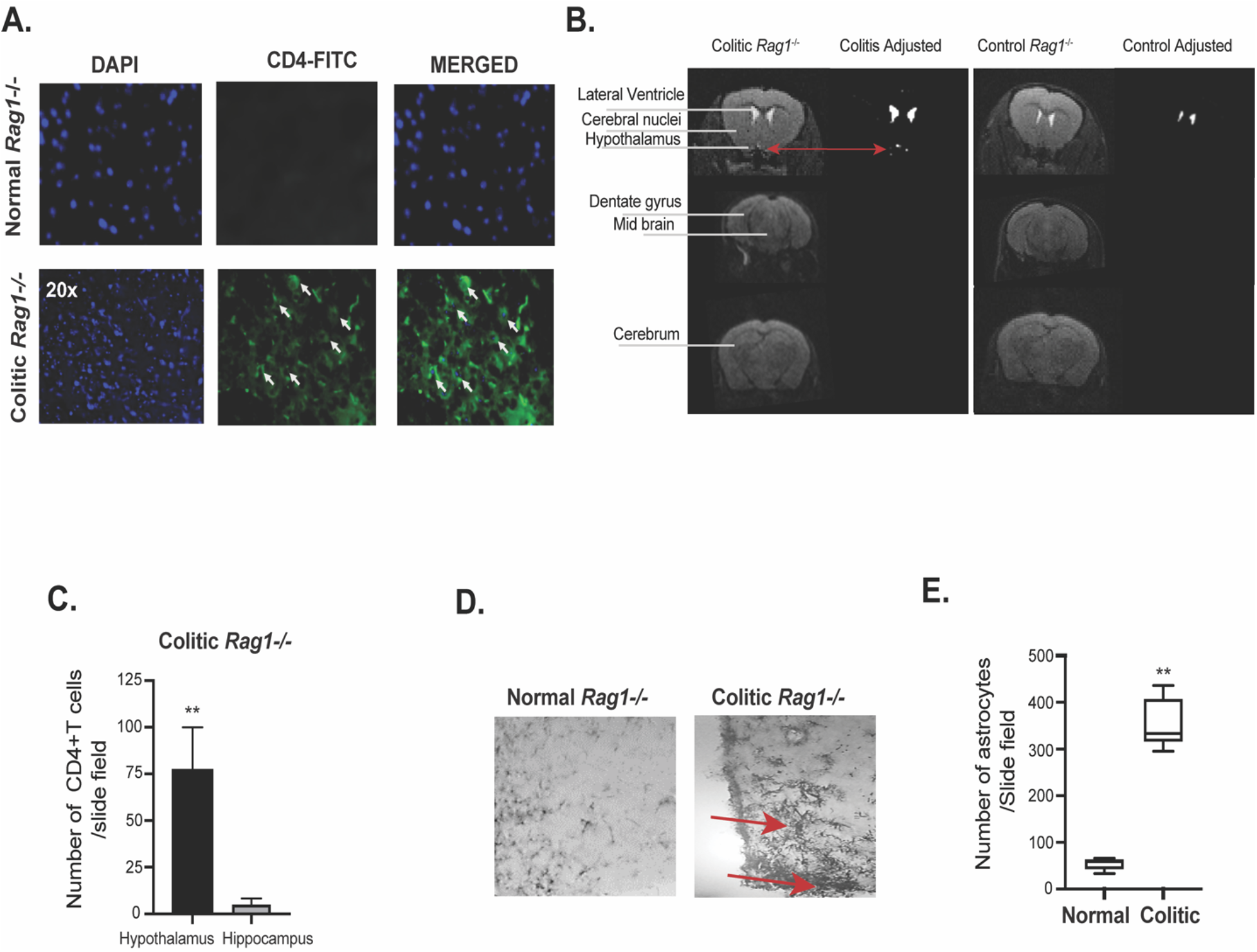
Evidence of brain inflammation in the hypothalamus of colitic mice accompanied with astrogliosis. **(A)** Immunofluorescence staining of CD4^+^ T cells in coronal brain sections of CD45RB^hi^ CD4^+^T cells transferred colitic *Rag1*^-/-^ mice and control un-transferred *Rag1*^-/-^ mice showing the presence of CD4^+^T cells. (**B**) Diffusion weighted MRI showing a hyper intense signal in the hypothalamus area of the brain in CD45RB^hi^ CD4^+^T cells transferred colitic *Rag1*^-/-^ mice at 10wk post-transfer compared to normal untransferred *Rag1*^*-/-*^ control. **(C)** Bar diagrams showing difference in the number of CD4^+^T cells between hypothalamic and hippocampal region of the brain of colitic mice at 8 wk post transfer. **(D, E)** Photo micrograph showing pronounced accumulation of astrocytes at the posterior hypothalamic paraventricular nucleus of the colitic mice at 20x magnification by GFAP staining of brain cross sections. Comparison of number of astrocytes in the hypothalamus showed ∼6 folds difference between colitis and the control mice. Data are shown as mean ± SEM. Data are representative of 2 independent experiments (C, E). *P* values, two-tailed unpaired Student *t*-test. **p<0.0001.

### Result 4. CNS-infiltrating CD4^+^T cell predominantly express RORγt and produce IL-17 that mirror proinflammatory colonic CD4^+^T cells

Both Th17 and Th1 cells are highly pathogenic effector cells in colitis and are comparable in their ability to induce colitis (46). Moreover, Th17 cells can cross blood brain barrier (BBB) more effectively than Th1 cells indicating that Th17 cells are more effective inducers of neuroinflammation (30). It has also been proposed that Th17 effector cells preferentially target astrocytes to promote neuroinflammation (31). Due to the presence of hypothalamic astrogliosis that corelated with the accumulation of CD4^+^T cells in the brain of colitic mice, we analyzed the brain-infiltrating CD4^+^T cells for expression of Th1, Th17 and Treg-specific TFs-T-bet, RORγt and FoxP3, respectively. CD4^+^T cells retrieved from colonic lamina propria (LP) of colitic *Rag1*^- /-^ mice expressed high level of RORγt (>50%) with moderate level of T-bet expression (<5%), indicating that RORγt-expressing CD4^+^T cells were the overwhelmingly dominant population in the colon (**Fig.4A**). Strikingly, similar to colonic CD4^+^T cells, brain-derived CD4^+^T cells of colitic mice also expressed very high level of RORγt (>20%). However, CD4^+^ T cells examined from all other organs didn’t express RORγt (**Fig. 4B, Fig. S3**). T-bet was only modestly expressed in the colon and brain CD4^+^T cells (2-4%) that was lowly expressed in all the other organs examined. However, FoxP3 expression was significantly higher in the colon and spleen and was nearly undetectable (<2%) in other organs including in brain CD4^+^T cells (**Fig.4 A, B**). This indicated that FoxP3^+^ Tregs were unable to migrate to the brain. Additionally, CD4^+^T cells of all other organs like spleen, liver, lung, peripheral lymph nodes (PLN) showed very low level of RORγt and T-bet expression (**Fig.4B** **& S3**), indicating that only CD4^+^T cells in the colonic LP and brain of colitic mice express high level of RORγt and moderate level of T-bet. As RORγt and T-bet are the master transcription factors of Th17 and Th1 cells respectively that induce production of proinflammatory cytokines IL-17 and IFNγ, we analyzed their expressions in different organs of colitic mice. In both IBD and MS, both Th17 cells and IFNγ co-producing ‘Th1 like’ Th17 cells are highly pathogenic (46, 47). While, frequency of IL-17 single positive and IL-17/IFNγ-double positive cells (∼5%) were highly induced in colonic CD4^+^T cells. Similar to colon, there was a significant population of IL-17 single producers (>40%) and IL-17A/IFNγ double producers in the brain (>12%) (**Fig. 4 C, D)**. However, frequency of IFNγ-single positive CD4^+^T cells were significantly lower in brain of colitic mice compared to the colon. As the frequency of pathogenic IL-17A/IFNγ co-expressing CD4^+^T cells was higher in the brain compared to the colon, it indicated that a significant proportion of brain infiltrating Th17 cells transition to pathogenic IFNγ producing Th17 cells-a process that can occur independent of T-bet expression (47). In contrast, IL-17 and IFNγ expressions were undetectable from all other organs examined in colitic *Rag1*^-/-^ mice (**Fig. 4 C, D)**. Together, our findings show that during colitis, Th17 and Th1-like Th17 cells are the dominant helper CD4^+^T cell population that are predominantly localized in the brain and in the colon, suggesting that proinflammatory characteristics of the LP-derived colonic CD4^+^T cells are represented in the brain-infiltrating CD4^+^T cells but absent from other major organs.

**Figure 4:**
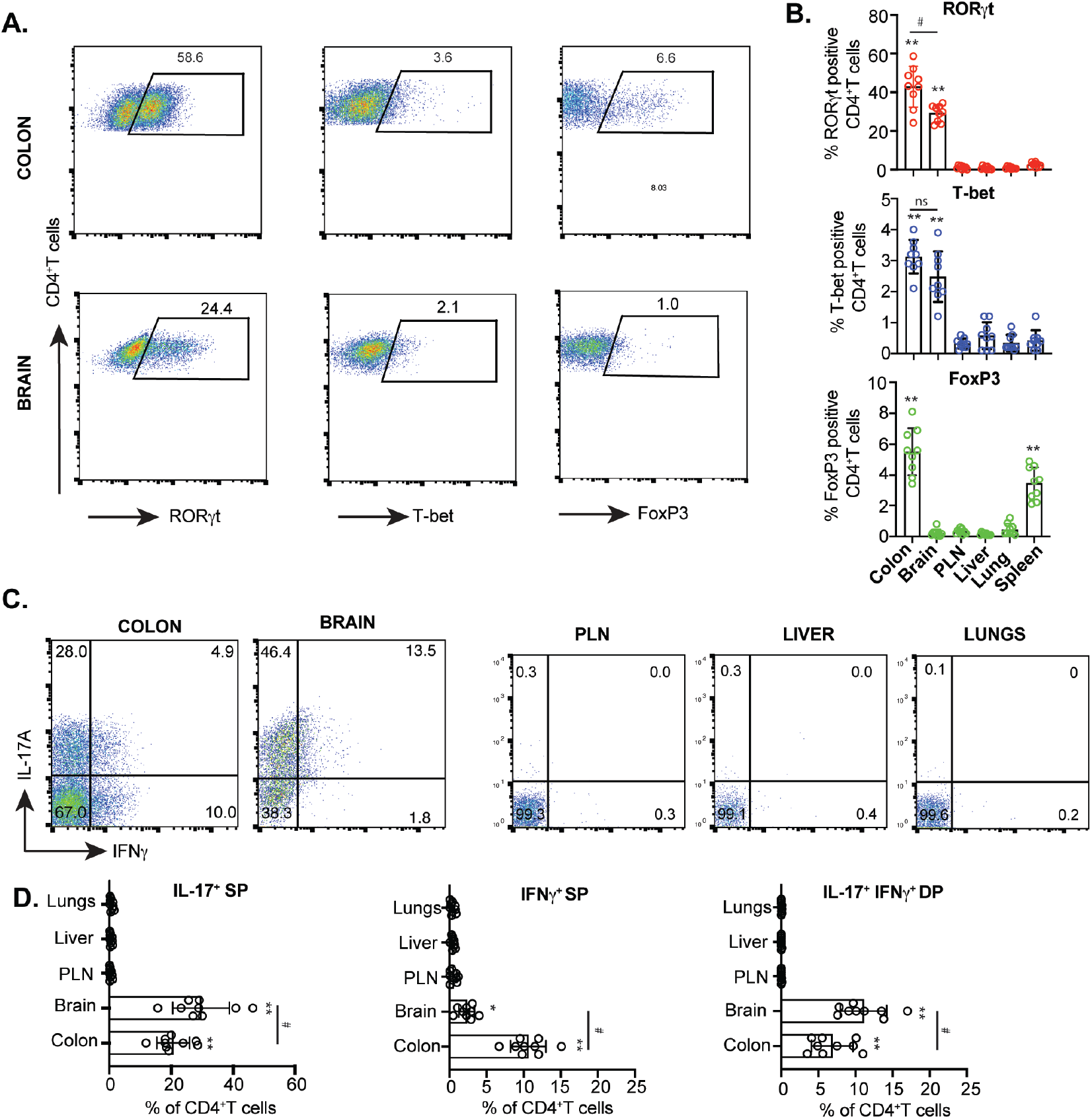
Both brain and colonic CD4^+^T cells express RORγt and IL-17 that are absent from CD4^+^T cells of other organs of colitic mice. **(A)** Representative FACS plots showing expressions of RORγt, T-bet and FoxP3 in live CD4^+^T cells retrieved from the brain and colonic LP of colitic *Rag1*^-/-^ recipient group at 8 wk post CD45RB^hi^ CD4^+^T cell transfer. **(B)** Bar diagrams showing percent RORγt, T-bet and FoxP3 expressing CD4^+^T cells retrieved from various indicated organs (n=9). **(C)** Representative FACS plots showing IL-17A and IFNγ expressions from CD4^+^T cells retrieved from the brain and colonic LP and other indicated organs of CD45RB^hi^ CD4^+^T cells recipient *Rag1*^-/-^ mice at 8 wk post transfer. **(D)** Bar diagrams showing percent IL-17A and IFNγ expressing CD4^+^T cells retrieved from various indicated organs (n=9). Data are shown as mean ± SEM. Data are representative of 3 independent experiments. *P* values, one-way ANOVA followed by Tukey’s post hoc test (B, D); #p<0.01; *p<0.005;**p<0.0001.

### Result 5. RORγt-deficiency impairs CD4^+^T cells infiltration into the brain of colitic mice

As the brain-infiltrating CD4^+^T cells overwhelmingly expressed RORγt, we investigated whether RORγt-deficient CD4^+^T cells can selectively infiltrate into the CNS during colitis. In support of a previous finding, we found that adoptive transfer of RORγt^-/-^ CD45RB^hi^ CD4^+^T cells transferred to *Rag1*^-/-^ mice caused mild colitis with significantly lower inflammation score (**Fig. 5A, B**). RORγt is essential for induction of IL-17A from CD4^+^T cells. We found that IL-17A expression from RORγt-deficient colonic CD4^+^T was nearly absent in *Rag*1^-/-^ recipient mice (**Fig. 5 C, D**). However, compared to WT colonic CD4^+^ T cells from reconstituted *Rag1*^-/-^ mice, IFNγ was strongly induced from colonic RORγt-deficient CD4^+^ T cells isolated from *Rag1*^*-/-*^ mice that received RORγt-deficient naïve CD4^+^T cells. This supports a previous observation that both IL-17A single producers and IFN-γ/IL-17A co-producers are more pathogenic and cause greater colonic epithelial damage than IFNγ single producers (46). In contrast, the recovered CD4^+^ T cells from RORγt-deficient naïve T cell recipient *Rag1*^*-/-*^ mice brain didn’t express either IL-17A or IFNγ. RORγt-deficient naïve T cells not only caused mild colitis, there was significantly lower number of CD4^+^T cells infiltration (<10-fold) in the brain of *Rag1*^-/-^ recipient mice as detected both at 4 and 8 wk post adoptive transfer (**Fig. 5 E, F**). Moreover, RORγt-deficient CD4^+^T cells in the brain did not express either IL-17A or IFNγ (**Fig. 5 G**). In contrast to the brain, number of colonic CD4^+^T cells did not vary significantly in the colon or other organs like spleen, lungs, PLN and liver between WT and RORγt-deficient CD4^+^ T cells recipient *Rag1*^-/-^ groups (**Fig. 5 H**). Together, these data suggest that infiltration of CD4^+^T cells in the brain is directly proportional to colitis severity and colonic RORγt-expressing CD4^+^T cells are essential for their homing from the colon to the brain during colitis.

**Figure 5:**
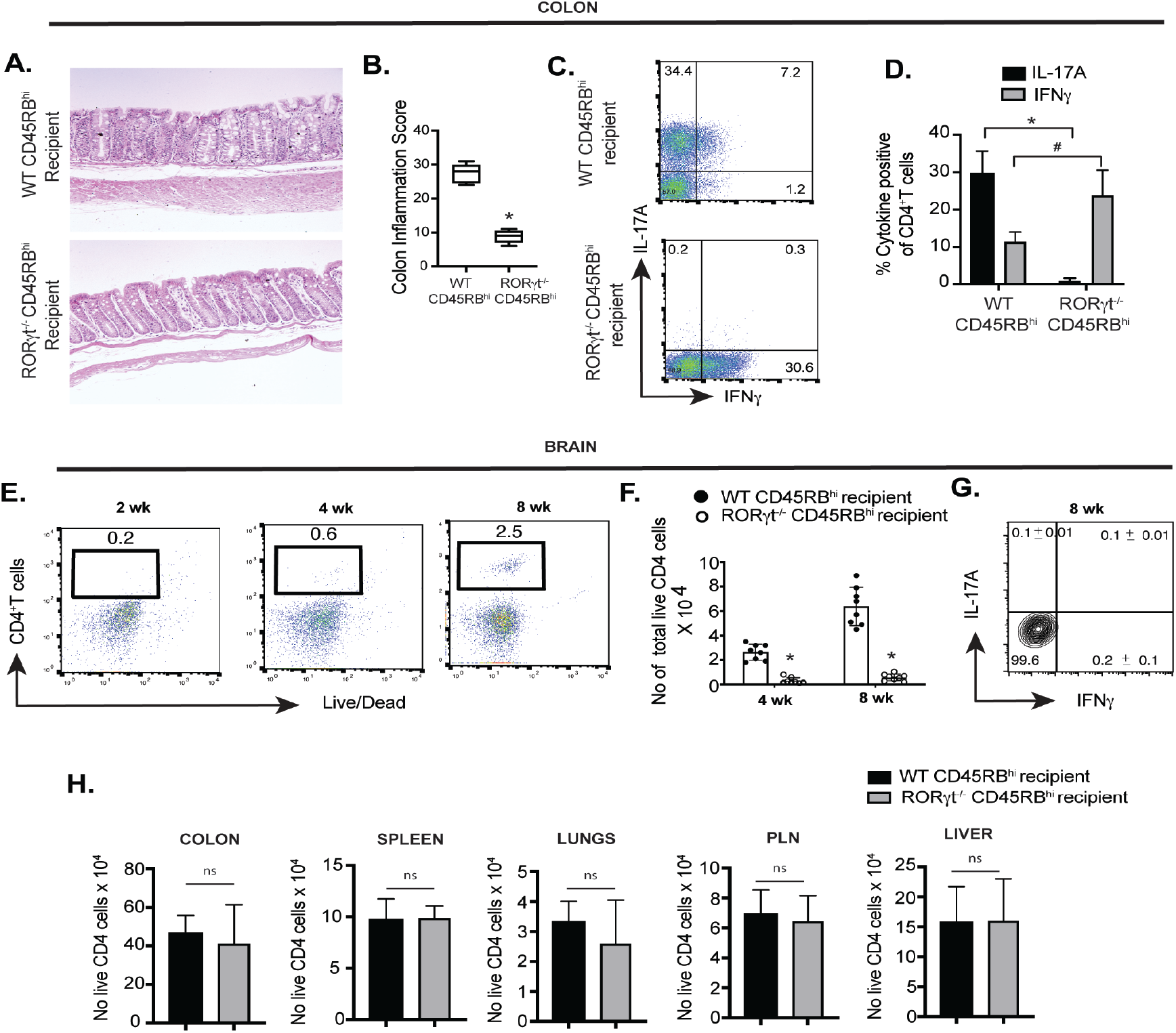
RORγt-deficient CD4^+^T cells are impaired in their ability to infiltrate into the brain. **(A, B)** Representative histopathology of H&E-stained colon sections along with inflammation score from *Rag1*^-/-^ mice adoptively transferred with either WT or RORγt-deficient naïve CD4^+^T cells 8 wk post. **(C)** Representative FACS plots showing expression of IL-17A and IFNγ from colonic CD4^+^T cells from colitic *Rag1*^-/-^ recipients at 8 wk post adoptive naïve CD4^+^ T cell transfer (n=8). **(D)** Bar diagram showing difference in percent IL-17 and IFN-γ expressing CD4^+^T cells isolated from colons of two groups of *Rag1*^-/-^ recipients that received either WT and RORγt^-/-^naïve CD4^+^ T cells. **(E)** Representative FACS plots showing frequency of CD4^+^T cells retrieved from the brain of *Rag1*^-/-^ mice adoptively transferred with RORγt-deficient naïve CD4^+^T cells. **(F)** Bar diagrams representing comparative analysis of total number of live CD4^+^T cells retrieved from brain of two groups of *Rag1*^-/-^ recipients (n=8). **(G)** Representative FACS plots showing expression of IL-17A and IFNγ from CD4^+^T cells retrieved from brain of *Rag1*^-/-^ mice that received RORγt-deficient naïve CD4^+^T cells. **(H)** Bar diagrams representing comparative analysis of total number of live CD4^+^T cells retrieved from various indicated organs of two indicated groups of *Rag1*^-/-^ recipients (n=6). Data are shown as mean ± SEM. Data are representative of at least two or more independent experiments. *P* values, two-tailed unpaired Student *t*-test; #p<0.001; *p<0.0001; n.s.=not significant.

### Result 6. Adoptive transfer of RORγt-deficient naïve CD4^+^T cells doesn’t cause brain inflammation and neurobehavioral disorders

To directly correlate absence of colitis severity and low number of brain-infiltrating CD4^+^T cells observed in RORγt^-/-^ naïve CD4^+^ T cells recipient *Rag1*^-/-^ mice, we initially compared extent of astrogliosis between WT and RORγt^-/-^ reconstituted groups. Accumulation of astrocytes in the hypothalamic region of the brain was significantly lower in *Rag1*^-/-^ recipient groups that received RORγt^-/-^ naïve CD4^+^T cells compared to WT (**Fig. 6A**). Moreover, the RORγt^-/-^ CD45RB^hi^ recipient *Rag1*^-/-^ mice showed no neurobehavioral abnormality compared to age-matched un-transferred *Rag1*^-/-^ mice with comparable behaviors in the EPM, sociability and light-dark tests, indicating absence of anxiodepressive-like behavior (**Fig. 6B**). Accordingly, RORγt-expressing Th17 cells that are critical for aggravating colitis severity are also crucial for causing brain inflammation and neurobehavioral disorders thereby linking colitis severity to neuroinflammation.

**Figure 6:**
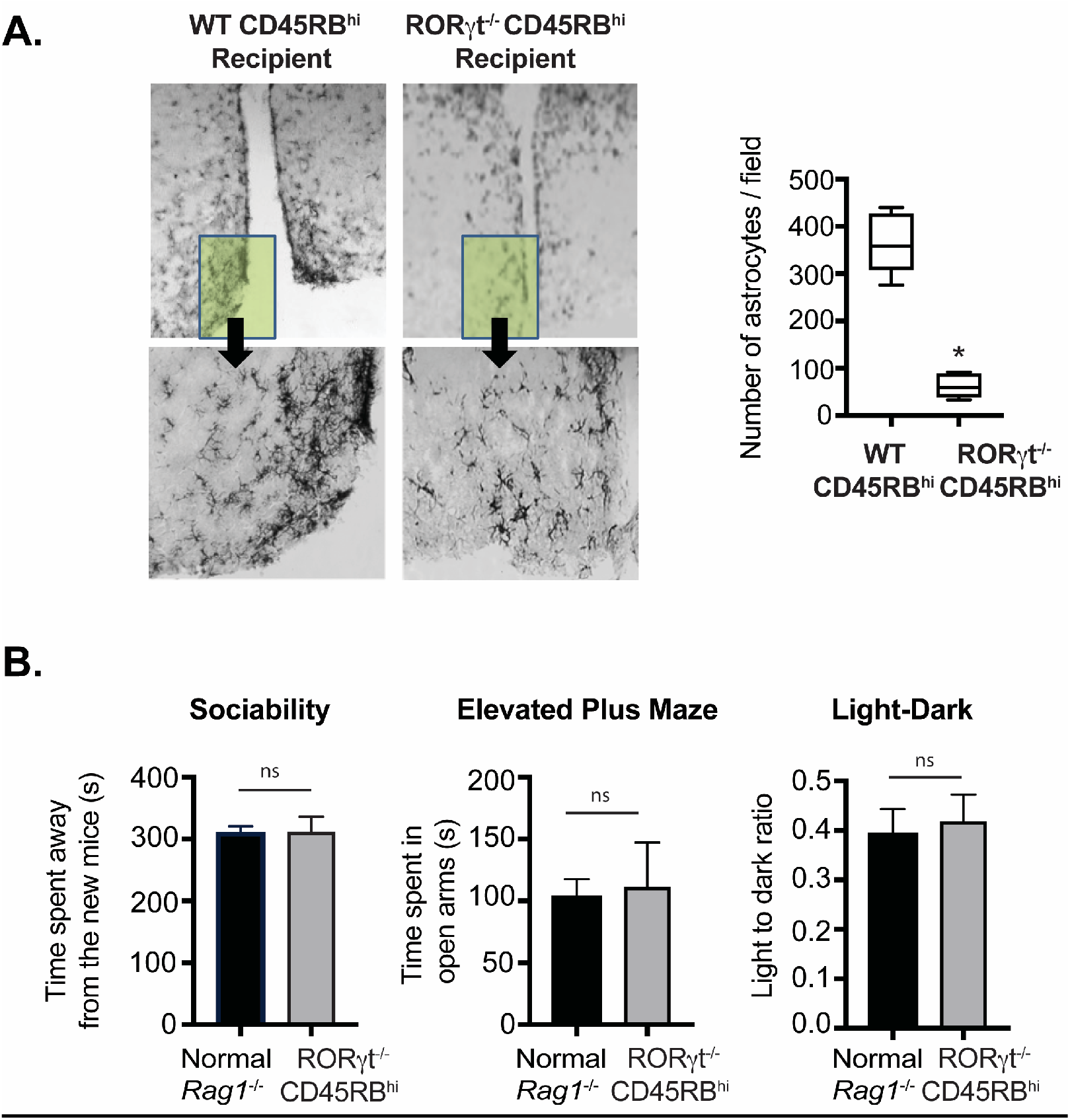
Adoptive transfer with RORγt-deficient CD4^+^T cells doesn’t cause brain inflammation and neurobehavioral disorders. **(A)** Photo micrograph at 20x magnification showing pronounced astrocytes accumulation by GFAP staining of coronal brain cross sections at hypothalamic region from two groups of *Rag1*^-/-^ recipients transferred with either WT and RORγt^-/-^CD45RB^hi^ CD4^+^ T cells. **(B)** Analysis of comparative neurobehavioral study by Sociability, Elevated Plus Maze and Light Dark test between indicated *Rag1*^-/-^ recipients that received RORγt^-/-^ naive CD4^+^ T cells and un-transferred *Rag1*^-/-^ groups. Data are shown as mean ± SEM. Data are representative of two to three independent experiments (4-5 mice /group). *P* values, two-tailed unpaired Student *t*-test (* p<0.0005); n.s.= not significant.

## Discussion

IBD patients suffer from several neuropathological comorbities (1-9), but the mechanisms behind these neurological manifestations during IBD remain largely unclear. Despite a growing field of research on the gut-brain axis, little is known about the mechanism by which gut pathology affects the brain. Our study found an intriguing mechanism that may directly link chronic intestinal inflammation to brain inflammation. In this study, we identified a cellular mechanism by which pathogenic RORγt-expressing CD4^+^T cells are correlated with the pathogenesis of colitis-driven neuropathology. These findings may help to explain how gut pathology can directly influence brain inflammation leading to neurological dysfunctions in IBD patients.

The brain-gut axis is a complex bi-directional system comprised of multiple connections between the nervous system and the gastrointestinal tract. It is believed that in IBD patients, psychological stress via the HPA axis increases intestinal motility and permeability leading to a local inflammatory response in the GI tract (20, 21). However, there is considerable debate on the “cause and effect” phenomenon between IBD and neurological comorbidity (10, 20). Evidence from preclinical studies suggests that intestinal inflammation can influence psychological behavior and brain activity (20). Studies of animal models with colonic inflammation have shown region-specific changes in the CNS that correlated with expressions of inflammatory genes in the brain (23-25). In support of this, a study has shown that CD3^+^CD4^+^T lymphocytes accumulate in a granuloma-like lesions in the brain of an IBD patient, suggesting that CD4^+^T lymphocytes infiltrate the brain during IBD and cause inflammation (48).

Classically, the brain and its associated structures have been considered to be immune-privileged (28). However, advances over the past decade have led to a reassessment of this assumption. New studies demonstrate the roles of a broad range of immune cells, including innate lymphoid cells and T cells, in regulating inflammation and neurological diseases in the brain (28). Researchers recently discovered infiltration of T cells in the aging brain (49). Intriguingly, pathogenic Th17 subset of CD4^+^T cells found in the brain of multiple sclerosis (MS) patients is also found in the gastrointestinal lesions of IBD patients making Th17 cells a common therapeutic target in both of these forms of autoimmune diseases(50-52). Th17 cells have been implicated in selective infiltration of the BBB and in causing neuroinflammation through the triggering of astrocyte activation (30-32). Several neurological disorders associated with cognitive impairment have been linked to Th17 cells, including neurovascular disorders and neurodegenerative diseases. Besides MS, Th17 cells are linked to various brain diseases, including demyelination, dementia, ischemic brain injury, Parkinson’s disease and Alzheimer’s disease (11, 14, 15, 32). However, little is known about the mechanism by which Th17 cells enter the brain to induce brain inflammation that might include either direct tissue destructive effects of IL-17 and IL-17/IFNγ co-producers on brain cells or indirect effects mediated through neurovascular dysfunction. A study has shown that due to the presence of IL-17R on endothelial cells (EC) of the BBB, Th17 cells transmigrate through BBB with greater efficiency than Th1 cells to promote CNS inflammation during MS (30). Interestingly, IBD patients also suffer from MS-like demyelinating conditions (11, 13, 15, 53). Therefore, it is likely that pathogenic CD4^+^Th17 cells that drive gut inflammation also contribute to inflammation of the brain during IBD.

Our study demonstrated that pathogenic Th17 cells, marked by the expression of their master TF RORγt, accumulate in the brain during colitis and cause brain inflammation. In view of the evidence suggesting that pathogenic CD4^+^Th17 cells traffic to the brain of colitic mice and trigger inflammation in the hypothalamus, we believe that neuropathological comorbidity in IBD patients may involve inflammation of the hypothalamus that is one of the major regions regulating anxiety and depression (17-19). It is noteworthy that brain is usually well protected from uncontrolled influx of molecules from the periphery due to the restriction imposed by the BBB, a physical seal of cells lining the blood vessel walls. However, the BBB around the hypothalamus that is located at the base of the brain, is a notable exception to this rule as this region is surrounded by “leaky” blood vessels that might readily allow entry of the CD4^+^T cells into the CNS (54, 55). Despite our observation that lymphocytes accumulation in the hypothalamus, it is quite possible that the invading CD4^+^T cells might eventually home to other regions of the brain at later stages of the disease and cause additional inflammatory changes. However, due to the limited time duration of the progress of the disease in the mouse model of chronic colitis, it’s difficult to validate the subsequent localization of CD4^+^T cells in other parts of the brain.

We have not demonstrated how RORγt-expressing Th17 cells selectively enter the brain during colitis. It is possible that RORγt plays an intrinsic role in either regulating the expression of specific adhesion molecule on Th17 cells that interacts with the ECs of BBB or by interacting with IL-17R of the ECs via the production of IL-17 to facilitate their attachment and migration through the BBB. Additionally, due to the inherent plasticity of Th17 cells, it is possible that they gradually become more pathogenic by increasingly transitioning to Th1-like IFNγ co-producing Th17 cells at later time points. Why the infiltrating Th17 cells continue to actively produce the proinflammatory cytokines after their infiltration in the brain should be an area of active investigation to determine if specific T cell clones are enriched in the brain by comparing their heterogeneity with the TCR clones of the colon during colitis. Identification of Th17 subsets with overlapping TCR repertoire will suggest that specific microbial antigens in colon trigger aberrant T cell response leading to their expansion and subsequent migration to the brain where they cross-react with overlapping brain antigens leading to their further expansion and retention of inflammatory potential.

This study suggests that the gut-brain immune axis as one of the principal corridors by which IBD drives the development of brain inflammation and behavioral neuropathology via migration of proinflammatory Th17 cells. We believe that the identification of specific molecules on Th17 cells that enable their infiltration into the brain by interacting with cognate receptors with the ECs of the BBB could lead to a novel therapeutic intervention that might be able to enhance the prognosis of both colitis and colitis-associated neuropathology.

## Materials and Methods

### Mice and Reagents

The following mice strains used were purchased from Jackson Laboratory: C57BL/6J (B6), *Rag1*^*-/-*^, B6(Cg)-*Rorc*^*tm3Litt*^/J. For all the experiment, 6-10 weeks old mice were used. Antibodies were purchased from either eBioscience, BD Biosciences or Fisher scientific for example-CD3, CD4, CD25, CD45RB, T-bet (eBio4B10), FoxP3 (FJK-16S), RORγt (AFKJS-9), IL-17A (TC11-18H10 or eBio17B7), IFNγ (XMG1.2).

### Chronic Colitis Induction

For adoptive T cell transfer model, CD25^neg^CD45RB^hi^ CD4^+^T cells (4 × 10^5^/mouse) from either *C57BL/6 or Rorc*^*fl/fl*^ *Cd4-Cre* mice were injected through intra peritoneal (i.p.) route in age and sex-matched *Rag1*^*-/-*^ mice (8-10 wk old, males or females) for colitis induction and recipient mice were monitored for 12 wk. For DSS-model of chronic colitis induction, mice were treated with 3 cycles of 2.5% Dextran Sulfate Sodium (Colitis Grade, Mol. wt. 36,000-50,0000; MP Biomedicals) dissolved in drinking water for 7 days and in between DSS-treatment, mice were kept in normal drinking water for 10-12 days (56).

### Isolation of CD4^+^ T cells from Brain, Colon and other organs and Antibody staining

For brain lymphocyte isolation, brain was isolated after intracardiac perfusion with PBS containing heparin to remove the blood from brain capillaries. Extracted brain tissue were cut into small pieces and incubated in RPMI containing collagenase IV (1 mg/ml, Sigma-Aldrich), Dispase (0.5 mg/ml, Gibco, Invitrogen) and DNase I (0.25 mg/ml, Sigma-Aldrich). for 20 minutes at 37°C shaker incubator. Then the mixture was filtered using 70-µm nylon cell strainers to capture single cell suspension. Mononuclear cells were isolated from the interphase between two Percoll (GE-Healthcare) gradients 70% and 38%. Cells were washed twice in cold PBS containing 1% FCS and subsequently used for antibody staining. Lamina propria lymphocytes (LPL) were isolated as described previously (57). Briefly, the large intestine was removed, cleared of luminal contents and fat, cut into small pieces and washed in chilled HBSS without Ca2^+^ or Mg2^+^. Minced tissue pieces were incubated in presence of EDTA for 30 min and vortexed to remove epithelial cells, incubated in RPMI containing digestion mixture. LPL were collected after passing through 70-µm strainers. For intracellular cytokine staining, cells were stimulated with PMA (50 ng/ml; Sigma) and ionomycin (750 ng/ml; Calbiochem) for 4 h in presence of Golgi Plug (BD Pharmingen). For CD4^+^T cells isolation from spleen, liver and peripheral lymph nodes (PLN) tissues were grinded and filtered through 70-µm strainers. Single cell suspension of spleen and liver were treated with ACK-lysis buffer to remove red blood cells followed by washing in cold PBS-1%FCS. Lung, kidney tissues were chopped and incubated in digestion mixture as stated before for 1h at 37°C shaker incubator and passed through 40-µm strainers (58). For detection of intracellular cytokines and transcription factor, cells were fixed and permeabilized either in FoxP3 staining Buffer (eBioscience) or BD permeabilization buffer. In all cases, LIVE/DEAD Fixable Near-IR Dead Cell Stain (Invitrogen) was included prior to surface staining to exclude dead cells in flow-cytometric analyses.

### Diffusion weighted MRI

Diffusion-weighted mouse MRI was conducted with a Bruker Biospec 9.4 Tesla scanner running Paravision 5.1 software (Bruker Biospin, Billerica, MA). A Bruker 72mm volume coil and Doty 24mm surface coil (Doty Scientific Inc., Columbia, SC) were used for signal excitation and reception respectively. Mice were anesthetized with isoflurane gas and monitored with an MRI-compatible physiological monitoring system (SA Instruments Inc., Stony Brook, NY). Mice were imaged in prone position with bite and ear bars for head fixation, and heated water was circulated through the animal bed to maintain animal body temperature throughout the experiment. Diffusion-weighted imaging of the brain was accomplished with a 2D spin-echo sequence using parameters TR/TE=3500/30ms, ETL=1, 1 average, FOV=1.92×1.92cm, and matrix 96×96 for an in-plane resolution of 200um. Slice thickness was set at 1mm, 7 slices were imaged with diffusion b-values of 0, 300, and 800 s/mm^2. Maps of the apparent diffusion coefficient (ADC) were calculated using custom code written in MATLAB (The MathWorks, Inc., Natick, MA). All images were transferred to our local PCs and analyzed using MATLAB Image Processing Toolbox.

### Immunofluorescence staining of Brain Sections

Following the perfusion of brain with PBS, tissues were extracted and fixed in 4% formaldehyde overnight. Brains were embedded in paraffin block followed by cut into 5-15µm coronal sections. Slides were immersed in xylene for 5 min followed by immersion in a series of ethanol concentrations for 2 min each. Slides were then immersed in boiled citric acid (0.01M) for 10 min followed by cooling and several washes in PBS. To avoid non-specific binding, the brain sections were incubated with 3% serum for 1 hour followed by over-night incubation with primary antibodies (1:1000; Rat anti-mouse CD4 from BD Bioscience) in a humidified chamber at 4°C. Next day, slides were washed twice with PBS and stained with secondary antibodies (1:400; streptavidin conjugated goat anti-rat IgG-Alexa fluor 488 from Invitrogen) for 2-3 hr at room temperature in a humidified chamber followed by incubation in prolong gold antifade reagent with DAPI (Invitrogen).

### Histology of Colon

Colon tissue samples obtained from proximal, middle and distal portions of large intestines were fixed in 10% neutral buffered formalin, embedded in paraffin to prepare 5 µm sections and stained with Hematoxylin and Eosin. The tissue sections were examined and scored to evaluate tissue pathology, as previously described (59). In all scoring, the identity of specimens was concealed from the pathologist.

### GFAP staining for Astrogliosis

For astrocytes staining, mouse-brains preserved in 4% paraformaldehyde, were cut using a manual slicing machine to 10μm thickness followed by rinsing in TBS. To quench endogenous peroxidase, sections were incubated with 3% H2O2 for 5 min at RT followed by incubation in a blocking solution containing 3% goat serum. The primary antibody (mouse anti-GFAP from Sigma at 1:1000) diluted in 0.5M TBS-Triton-X-100 (0.5%, pH 7.6) buffer was added in humidified chamber on a shaker table in the dark overnight at RT. The slices were rinsed in TBS-T (5 min each) followed by the addition of secondary antibody (Goat anti-mouse IgG-biotin from Sigma, 1:500) and kept at RT for 2 hours in a humidified chamber. The tissue was rinsed for 3 times, in 20 ml TBS-T. A tertiary streptavidin-HRP antibody (a part of the Mouse ExtrAvidin® Peroxidase Staining Kit from Sigma at 1:1000) was added for 2 hr, followed by diaminobendizine (DAB) substrate reaction for 2-3 min. Finally, the tissue was rinsed to remove excess DAB and mounted.

### Luxol Fast Blue-PAS staining for Myelin

For analysis of myelin integrity, paraffin-embedded coronal brain sections were dipped in xylene twice for 5 mins each followed by 3 changes in absolute alcohol and rinsed in deionized water for 1 min. Deparaffinized brain sections were placed in 0.1% Luxol Fast Blue solution at 56 °C for 1 h in water bath followed by quick dip in 0.05% lithium carbonate solution and then in 70% alcohol. For counter stain, slides were immersed in 0.5% Periodic acid for 10 min followed by incubation in Schiff’s solution (Sigma) for 10 mins followed by washing in water and dehydrated sequentially in 70% alcohol and in absolute alcohol, cleared in xylene and mounted.

### Behavioral experiments

#### Light-dark box test

This test is used to asses anxiety-like behavior. The light/dark inclination assessment was executed by means of automated tracking device consisting of a video camera tracker connected to a PC (Med Associates, Inc, St Albans, VT). The test itself was done using a light-dark box made of plastic. Each mouse was positioned in the middle of a box subjected to normal illumination along with no illumination in the adjacent box and their movements were recorded for 10 mins. Measurements included the total time variation between the two boxes (light vs. dark) and number of entries to each compartment (60).

#### Elevated plus maze

This experiment was used to evaluate depression and anxiety-related behavior in rodents. The maze is composed of four wooden arms of equal length with two closed arms and two opened arms and a central area between the arms. Anxious mice tend to shelter in more secure area of the closed arms, while mice that do not show anxious behavior prefer to spend more time investigating the open arms. To perform the experiment each mouse was positioned in the center area and it was left to freely choose the area it wants to stay in or explore for 5 mins. The time spent in each area was calculated (61) (62).

#### Social interaction test

This test was performed to explore approach-avoidant behavior as an indication of anxiety-like behavior according to Crawley’s standard protocol. The test takes place in a rectangular box consisting of three chambers (63). In one of the chambers, an unknown control mouse was placed in a wired cup, while the other room was left empty. In this test, the mouse was given a chance between inspecting the unknown mouse placed in the wired cup or inspect the empty room. To ensure the accuracy of this test, the box was equally lit by general room lighting, and all mice were given time to acclimate for 30 mins before the test started. Habituation (adaptation) was performed for 10 min. To assess anxiety, mice were monitored for 10 min in the chamber. The time consumed by the mouse in each respective chamber along with the frequency of entries to each room was calculated.

#### Y-maze

Y-maze test was performed to assess short term memory for mice suffering from colitis. The test setup consists of “Y” shaped plastic corridor. The test was performed in two phases. In the first phase one of the arms was closed with a wooden door, while the mouse was allowed to explore the other two arms for 8 minutes. In the second phase of the test (8 minutes), the opened and the closed corridors forming the “Y” arms are exchanged. Mice that do not show signs of spatial memory deficits tend to remember which arms they had explored and thus will explore the newly opened arm (64).

#### Fear conditioning

This measures the mouse’s ability to recall incidents through the retrieval of information from short-term memory storage. In this test, each mouse was subjected to an aversive unconditioned stimulus (e.g., short electrical shock) associated with a sound of 90db for 30 secs. The conditioning of the sound and the electric shock was repeated twice on the first day of the experiments with 90 secs between the shock and the second. On the second day, the ability to retrieve the association was calculated by calculating the freezing time shown by the mice. In order to ensure the accuracy of the test, the mice were allowed to explore the fear condition chamber (Med Associates/Actimetrics chamber system) for 3 mins on the first day, while on the second day, mice were allowed to explore the cage for 5 mins to help retrieve short-term memory conditioning (62).

#### Novel Object recognition test

This test was performed to measure the mice’s ability to differentiate between objects that are ‘familiar’ versus ‘new’ objects. The medium of this experiment is a square plastic field with closed sides. The arena is divided into two sides, with an object placed on each side. In the first phase, the mouse was put in the arena and left to explore the two sides containing the two objects for 10 min. In the next phase of the experiment, one object was substituted with a novel item and mouse was left freely to explore the objects for 5 min. The duration of inspecting the novel object was compared to the time spent exploring the old object. Mice that do not have short term deficits will spend a longer time with novel objects(61).

#### Horizontal Ladder

This test was used to measure muscle coordination by the cerebellum. In the test, mice have to stroll across a corridor consisting of a horizontal ladder that ends with a covered area. Mice tend to prefer to stay in the covered areas given the ability to move. The spaces between each rung are paced at 1 cm, to make sure the mouse can coordinate its movement without miss stepping between the rungs. The number of missteps per each mouse was recorded, together with the total time to walk from one end of the ladder to the other.

### Statistics

All the statistical analysis was done by GraphPad Prism software. *P*-values were calculated by Pearson’s r test to measure linear correlation between two sets of data, two-tailed unpaired Student’s *t*-test, two-tailed paired Student’s *t*-test, one-way ANOVA followed by Tukey post-hoc test as described in figure legends. All *p*-values ≤ 0.05 was considered significant.

### Study Approval

Mouse studies conducted at UAB were approved by Institutional Animal Care and Use Committee and all relevant ethics regulations were followed.

## Author Contributions

MEM designed, planned and performed experiments; analyzed data; and wrote the manuscript. SB performed experiments; assisted with data interpretation and experimental design; edited the manuscript. AC, AU, JT performed experiments. TVG provided reagents and supervised experiments. AGS, MM assisted with data interpretation. JAB, DGS, supervised the study and edited the manuscript. RB planned and supervised the study; analyzed the data and manuscript writing.

## Notes

This study has been supported by a Career Development Award (RB) from the Crohn’s and Colitis Foundation of America (Identifier# 347717); and Start-up fund from UAB school of Medicine (RB); NIH grants MH079710 and MH116896 (JAB); T32NS061788 (A.U.); NIH (Yale/NIDA Neuroproteomics Center grant DA018343, AC); Alabama Udall Center, NS108675 (DGS).

**Disclosures: The authors have no financial conflicts of interest.**

### Competing Interest Statement

The authors have declared no competing interest.

