## Supplemental Figures 1-3 for "RORγt-Expressing Pathogenic CD4^+^T Cells Cause Brain Inflammation During Chronic Colitis"

<u>**SUPPLEMENTARY FIGURE 1**</u>

Supplementary Figure. 1

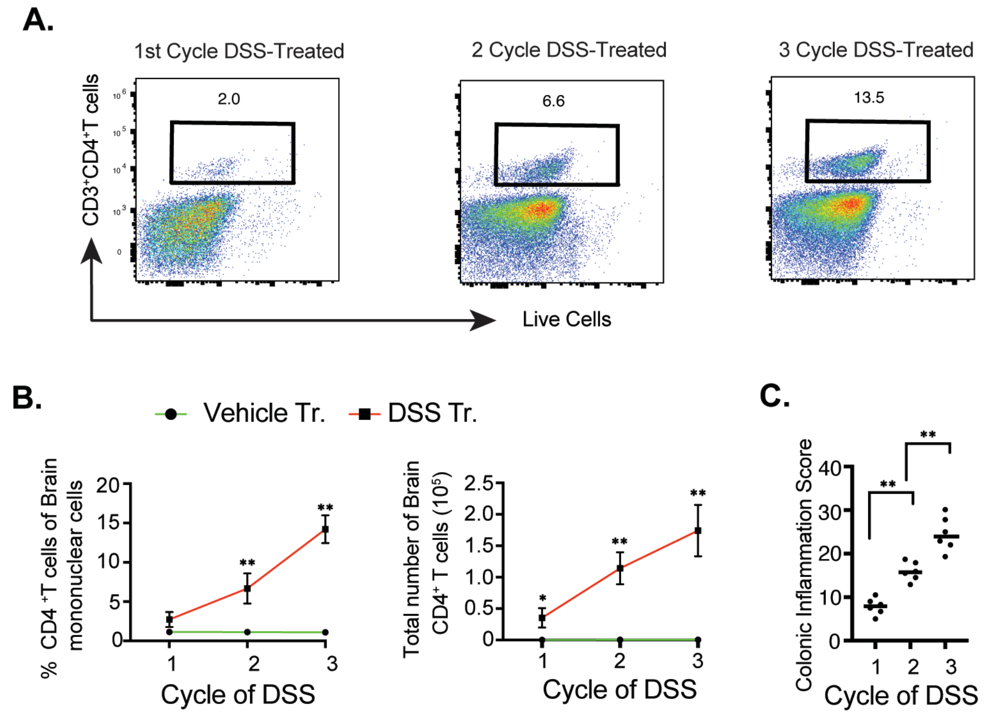


**Supplementary Figure S1**: **CD4⁺ T-cell infiltration in the brain during DSS-induced chronic colitis. (A)** C57BL/6 mice were subjected to 3 cycles of 2.5% DSS in drinking water for one week followed by 10-14 days in normal drinking water in between DSS-cycles for colitis induction. Brain infiltrating $CD4^+$T cell frequency was analyzed by flow cytometry after each cycle of DSS-treatment. **(B)** Kinetics of $CD4^+$ T cells infiltration in the brain of DSS-treated mice presented as percent (left) and total number (right) following each cycle of DSS-treatment. **(C)** Intestinal inflammation score of DSS-treated mice after each cycle of treatment. Data are shown as mean ± SEM (C). Data are representative of three independent experiments (6 mice /group). *P* values, one-way ANOVA followed by Tukey's post hoc test (B) and two-tailed paired Student *t*-test (C) * $p< 0.05$; ** $p< 0.001$.

### SUPPLEMENTARY FIGURE 2

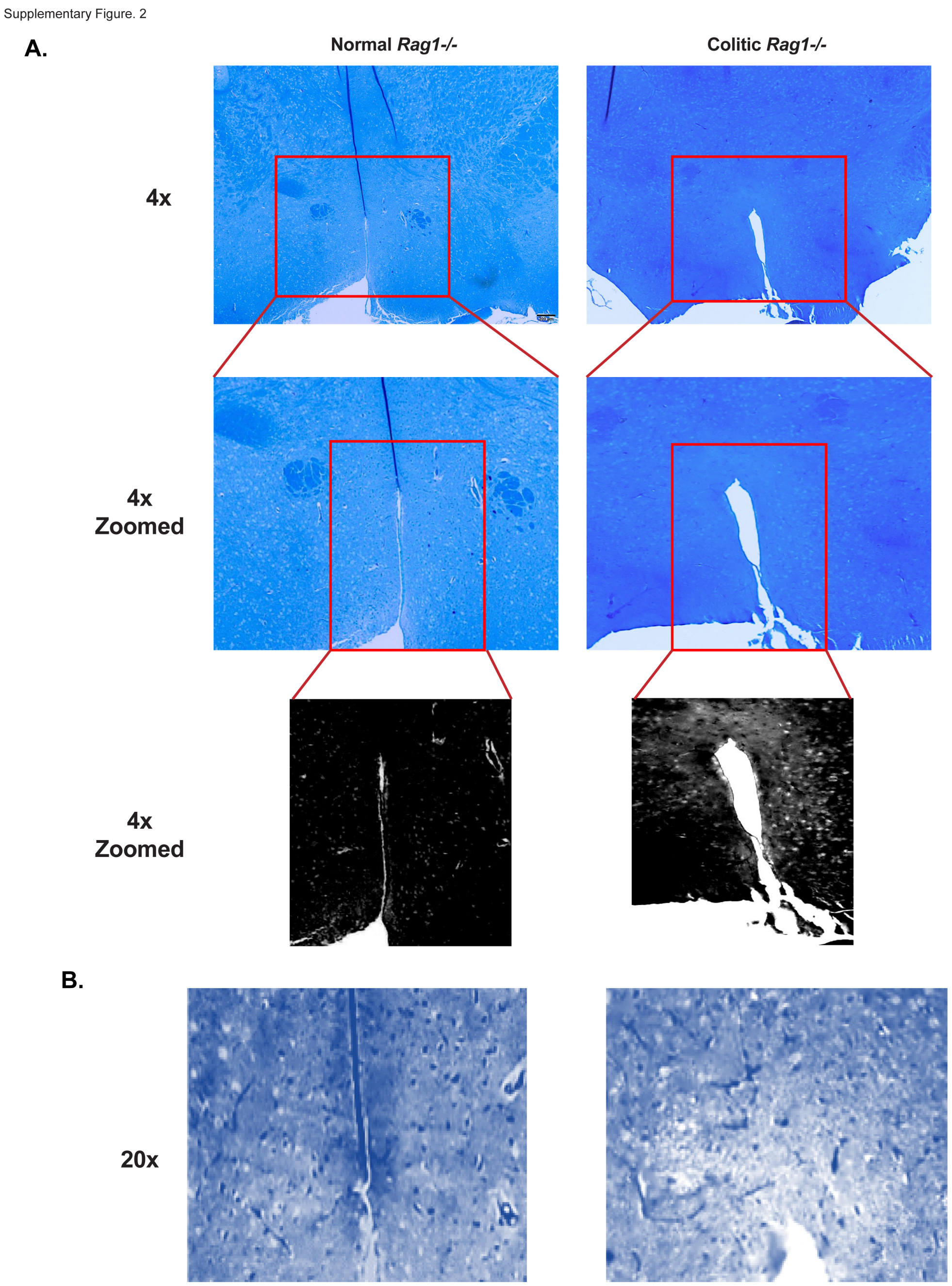


**Supplementary Figure S2**: **Evidence of demyelination in the hypothalamic region of colitic mice.** Luxol fast blue-PAS staining on the brain cross sections showing marked decrease in myelination in the hypothalamic region of CD45RB$^{hi}$ CD4$^{+}$T cells recipient *Rag1*$^{-/-}$ mice compared to untransferred *Rag1*$^{-/-}$ control mice at 4x or 4x zoomed magnification **(A)** and at 20x magnification **(B)**.

**SUPPLEMENTARY FIGURE 3**

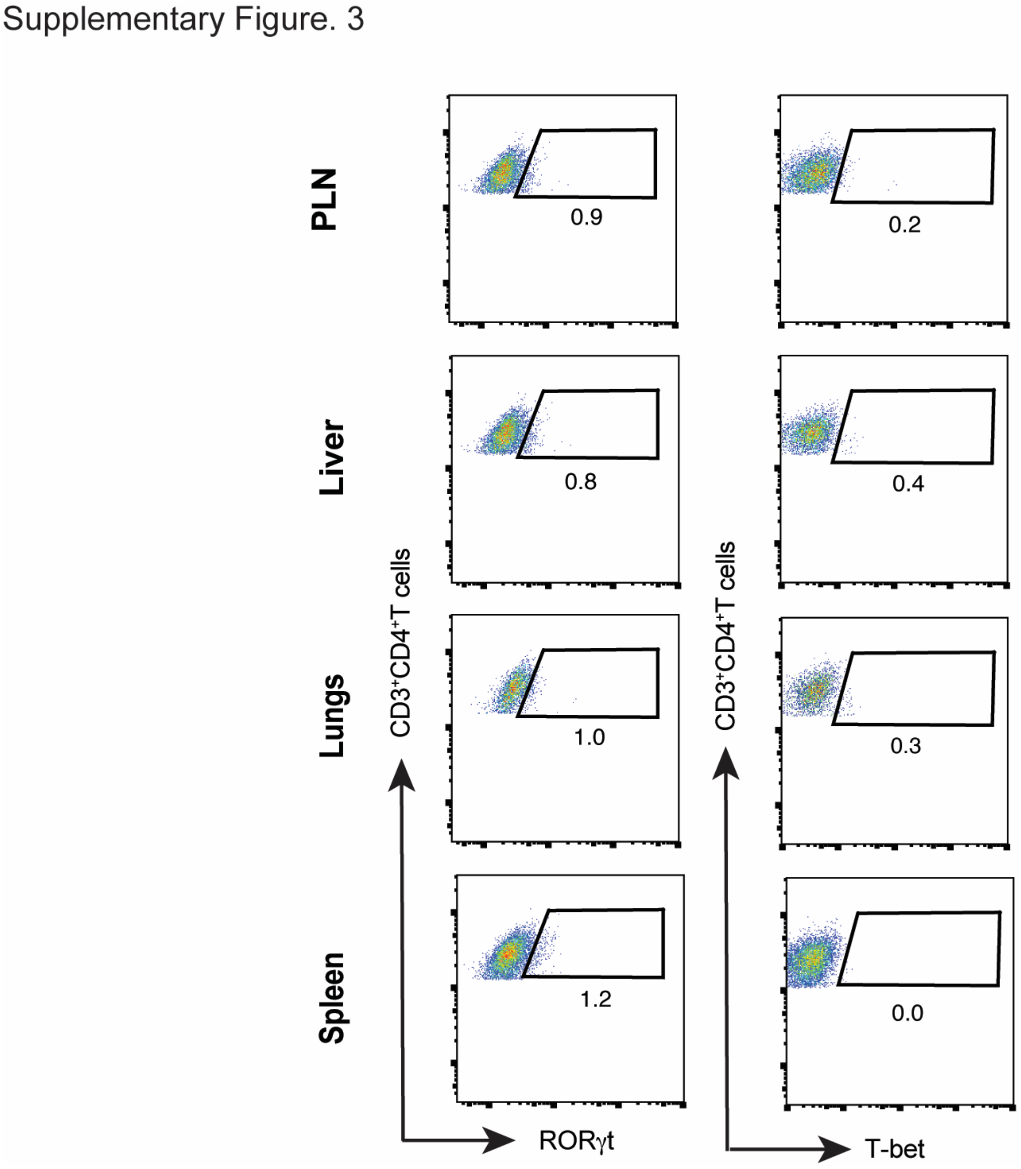


**Supplementary Figure S2**: **Expression of RORγt and T-bet from CD4$^+$T cells in colitic *Rag1$^{-/-}$* mice in the indicated organs.** Representative FACS plots showing RORγt and T-bet expressions in live CD4$^+$T cells retrieved from the PLN, Liver, Lungs and Spleen of colitic *Rag1$^{-/-}$* recipient group at 8 wk post CD45RB$^{hi}$ CD4$^+$T cell transfer.
